# Hunter-gatherer genomes reveal diverse demographic trajectories following the rise of farming in East Africa

**DOI:** 10.1101/517730

**Authors:** Shyamalika Gopalan, Richard E.W. Berl, Gillian Belbin, Christopher R. Gignoux, Marcus W. Feldman, Barry S. Hewlett, Brenna M. Henn

**Affiliations:** Department of Ecology and Evolution, Stony Brook University, Stony Brook, NY 11794, USA.; Department of Human Dimensions of Natural Resources, Colorado State University, Fort Collins, CO 80523, USA.; Icahn School of Medicine at Mount Sinai, New York, NY 10029, USA.; Department of Medicine, University of Colorado, Anschutz Medical Campus, Aurora, CO 80045, USA.; Department of Biology, Stanford University, Stanford, CA 94305, USA.; Department of Anthropology, Washington State University, Vancouver, WA 98686, USA.; Department of Anthropology, University of California, Davis, CA 95616, USA.; UC Davis Genome Center, University of California, Davis, CA 95616, USA.

## Abstract

A major outstanding question in human prehistory is the fate of hunting and gathering populations following the rise of agriculture and pastoralism. Genomic analysis of ancient and contemporary Europeans suggests that autochthonous groups were either absorbed into or replaced by expanding farmer populations. Many of the hunter-gatherer populations persisting today live in Africa, perhaps because agropastoral transitions occurred later on the continent. Here, we present the first genomic data from the Chabu, a relatively isolated and marginalized hunting-and-gathering group from the Southwestern Ethiopian highlands. The Chabu are a distinct genetic population that carry the highest levels of Southwestern Ethiopian ancestry of any extant population studied thus far. This ancestry has been in situ for at least 4,500 years. We show that the Chabu are undergoing a severe population bottleneck which began around 40 generations ago. We also study other Eastern African populations and demonstrate divergent patterns of historical population size change over the past 60 generations between even closely related groups. We argue that these patterns demonstrate that, unlike in Europe, Africans hunter-gatherers responded to agropastoralism with diverse strategies.

---

Since the beginning of the Holocene 12,000 years ago (ya), the dominant mode of human subsistence has shifted from hunting and gathering to agriculture through a process known as the Neolithic transition. Whether this transition occurred primarily through the mass movement of people from centers of domestication (demic diffusion) or through the cultural transmission of agricultural practices (cultural adoption) is still debated in archaeology, genetics and anthropology (*1*, *2*). As this transition largely concluded by 4,000 ya in Europe and Asia, there remains little direct evidence of on-the-ground interaction between hunter-gatherers and agriculturalist migrants. Theoretically, in the face of displacement and conflict over resources, hunter-gatherer populations might respond in a variety of ways: 1) intermarry with the migrant group and adopt their agropastoral subsistence practices (substantial genetic exchange); 2) adopt the subsistence practices of the migrants without intermarriage (limited genetic exchange); 3) reduce their geographic range or resource acquisition (leading to a decline in population size); 4) enter into an economic-symbolic exchange relationship with the migrant group; or 5) move to an ecological region that is marginal for pastoralism or agriculture (*3*, *4*). These are not mutually exclusive; the history of any particular hunter-gatherer group may involve multiple modes of response. In Europe, the current consensus is that early Near Eastern farmers facilitated the spread of agriculture, completely replacing groups in some areas, and in others – particularly in southern Europe – admixing with hunter-gatherers (*5*–*8*). However, the extent to which the Neolithization processes of Europe occurred on other continents remains unclear. In Africa, the genetic and cultural landscape has been significantly shaped by recent expansions of agriculture and pastoralism, such as the Bantu migration (*9*, *10*), but a lack of data precludes detailed characterization of these expansion events.

We sought to understand the demographic impact of the expanding Neolithic in Southwest Ethiopia, a region of high ethnic and linguistic diversity that is home to several of the world’s remaining hunter-gatherer groups, but has a relatively sparse archaeological record (*11*). We present new genomic data from the Chabu hunter-gatherers and their immediate neighbors, the Majangir and Shekkacho, which we analyze together with other East African populations. The Chabu people are poorly known even among anthropologists. They inhabit the highland forests that straddle the border between the Oromia Regional State, Gambella Regional State, and Southern Nations, Nationalities, and Peoples’ Region (SNNPR) (*12*). Recent estimates of their census size range between 1,700 and 2,500 individuals (*12*). The Chabu claim to be the original inhabitants of these forests, an assertion that their Majangir and Shekkacho neighbors generally support. The Chabu are thought to be a linguistic isolate, while the Majangir are Nilo-Saharan Surmic-speakers who practice small-scale cultivation and limited hunting and gathering and the Shekkacho are Afro-Asiatic Omotic-speakers who practice intensive agriculture (*13–15*).

We investigate two alternative hypotheses regarding the origins of the Chabu. The first is that the Chabu are the descendants of an earlier population that occupied Southwest Ethiopia prior to encroachment by agriculturalists. Archaeological research from Ajilak, roughly 100 km west of the Chabu forest and 500 m lower in elevation, strongly suggests that hunter-gatherers were present in the area as late as 800-1,000 years ago (*16*). The presence of iron tools from Ajilak indicates that foraging groups interacted with lowland Sudanese pastoralist populations in the past. Based on archaeological and ethnographic evidence, it has previously been suggested that the Chabu, the Majangir, and Koman-speakers could be the living descendants of the Ajilak hunter-gatherers (*16*). A competing hypothesis is that the Chabu were previously an agricultural or agropastoral group that transitioned to foraging in order to exploit an unused ecological niche in the forest highlands. This hypothesis implies close relatedness between the Chabu and another Ethiopian or Sudanese population, from which they would have diverged relatively recently. Although a transition to hunting and gathering from other subsistence modes is presumed to be uncommon historically, it has occurred more than once (*17*, *18*). In Eastern Africa specifically, some primarily pastoralist groups are known to fluctuate between subsistence strategies (*19*, *20*).

In order to test whether the Chabu are descendants of earlier hunter-gatherer inhabitants, we performed unsupervised clustering of autosomal single nucleotide polymorphism (SNP) data from populations of Ethiopia, Sudan, South Sudan, and Kenya (Methods). Importantly, we also included genomic data from Mota, a 4,500 year old individual found in the nearby Gamo highlands who lived prior to any evidence of agriculture or pastoralism in the region (*21*). In the clustering algorithm, we varied ‘K’, the hypothesized number of ancestral source populations for the dataset, from 2 to 12. We focus on the pattern that emerges at K=5, and refer to these five genetic components as ‘Nilo-Saharan’, ‘Afro-Asiatic’, ‘Southwestern Ethiopian’, ‘Niger-Congo’, and ‘Near Eastern’ ancestries based on their frequencies across geographic space and language families (Fig. 1). The Chabu and their neighbors are primarily characterized by differences in frequency of the first three components. Nilo-Saharan ancestry is most concentrated in southern Sudan and South Sudan, overlapping slightly with western Ethiopia (i.e. at high frequency in the Nilo-Saharan-speaking populations of the region) (Fig. 1B). Afro-Asiatic ancestry is highest in northeast Ethiopia, Eritrea, and northeast Sudan, declining steadily towards the south and west (Fig. 1C). Southwestern Ethiopian ancestry is found in all language groups but is concentrated in the Southwestern Ethiopian highlands (Fig. 1D). Importantly, the ancient Mota genome carries a high proportion of this ancestry, suggesting that it has characterized this region for at least 4,500 years. Among the contemporary groups we studied, the Chabu carry the highest levels of Southwestern Ethiopian ancestry, as well as moderate levels of Nilo-Saharan ancestry (Fig. 1A). The Majangir and the Gumuz are characterized by the same components, each present at ~50%. The Chabu are distinct from Afro-Asiatic Ethiopian and Sudanese agropastoral populations, as well as from traditionally pastoral Nilotic speakers, who include the Anuak, Dinka, and Shilluk (Fig. 1A). Similarly, in principal component (PC) space, the Chabu anchor the second PC and are closest to Mota, the Majangir, and Gumuz (Fig. S1). Notably, the Majang and Gumuz are not readily distinguishable from each other in either K=5 ADMIXTURE plots or a biplot of the first two PCs. We conclude that the Chabu carry the major Southwestern Ethiopian genetic component also identified in Mota, and are distinct from agricultural and/or pastoralist groups. Therefore, they are likely to be direct descendants of ancient Southwest Ethiopian hunter-gatherer groups; secondary adoption of hunting-and-gathering is not supported.

**Figure 1.**
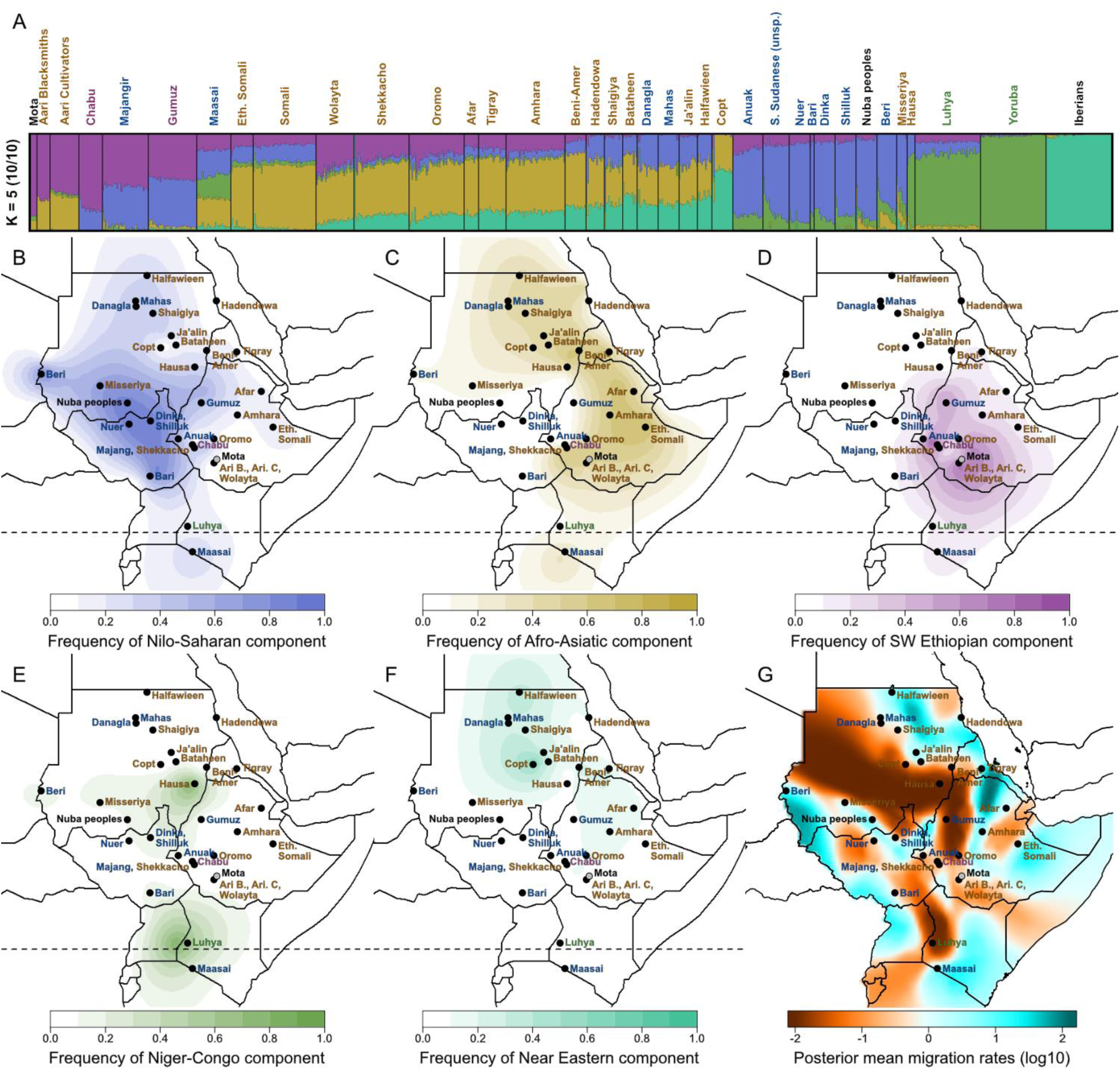
Global ancestry proportions of northeast African individuals, the Yoruba from Nigeria and Iberians from Spain, inferred using unsupervised clustering. A) Each color corresponds to a genetic component and each vertical bar represents one individual (Mota is plotted 5 times wider for visualization purposes). The population labels are colored according to linguistic affiliation, with green denoting Niger-Congo speakers, blue Nilo-Saharan speakers, yellow Afro-Asiatic speakers, and purple speakers of linguistic isolates. All ten replicates at K=5 ancestral components converged on the same overall pattern of partitioning the genetic variation. B-F) The geographic distributions of these components are depicted, with the intensity of the color corresponding to the mean population proportion of the respective ancestry. G) Effective migration surfaces, inferred using the rate of decay of genetic similarity across space, are depicted. Cool colors correspond to effective migration corridors, while warm colors correspond to effective migration barriers.

Despite carrying moderate to high levels of Southwestern Ethiopian ancestry, there are clear differences between the Chabu-Majangir-Gumuz and the Aari-Mota genetic profiles. At K=5, Mota and the Aari carry Afro-Asiatic ancestry but no Nilo-Saharan ancestry. In a biplot of the first two PCs, the Aari populations pull away from the Chabu-Majangir-Gumuz towards the Wolayta, the Afro-Asiatic group who are their nearest neighbors today (Fig S1). These ancestry differences may relate to differential gene flow with neighbors or may reflect ancient patterns of divergence.

Next, we modelled spatial population structure by using genetic data to estimate the effective migration surface (EEMS) (*22*). This analysis reveals corridors of and barriers to migration that closely correspond to the spatial distribution of ancestral components (Fig. 1G). Some migration barriers also correspond closely with major geographic features such as deserts, certain high elevation areas, and bodies of water. However, other major features, such as the Nubian Desert and Northeastern Ethiopian Highlands, do not appear be have been historical barriers to migration (Fig. S2). Furthermore, areas with low rates of migration tend to lie along the boundaries between the Nilo-Saharan, Afro-Asiatic, and Niger-Congo language families, while corridors of high gene flow lie within them (Fig. S2C). Together, these results emphasize the close association between geography and language in determining gene flow between groups (*23*). Furthermore, the Chabu lie directly in the center of a language contact area with negative effective migration rates, perhaps indicating relative isolation from neighboring groups.

During the last decade, the Chabu have faced mounting challenges to their survival. Their land claims are not recognized, and as a result their traditional forests are often violently co-opted for coffee plantations and other development (*12*). We sought to understand the effects of recent and historical isolation and demographic pressures faced by the Chabu by analyzing runs of homozygosity (RoH) (Methods). Compared to their neighbors, the Chabu carry more of their genome in RoH (Fig. S3). As the Chabu have no cultural tradition of close relative marriage and practice clan exogamy, this demonstrates an increased level of isolation relative to their neighbors (*12*). We compared the Chabu to other African hunter-gatherers, including the Biaka, Mbuti, and Hadza, as well as recent descendants of hunter-gatherer groups; these include the Majangir, Gumuz, Sandawe, and Aari. The former three groups are primarily small-scale farmers today, but ethnographic studies indicate that they regularly hunted and gathered in the recent past, while the genetic similarity of the Aari to Mota indicate that they are also recent descendants of hunter-gatherers (*14*, *21*, *24*–*27*). Compared to other Ethiopian populations that carry a large proportion of Southwest Ethiopian ancestry, the Chabu have the highest levels of total RoH (Fig. 2). However, the Hadza of Tanzania, who were previously shown to carry the highest proportion of their genomes in RoH (total RoH) among Africans, exceed the Chabu in this measure (Fig S4) (*26*).

**Figure 2.**
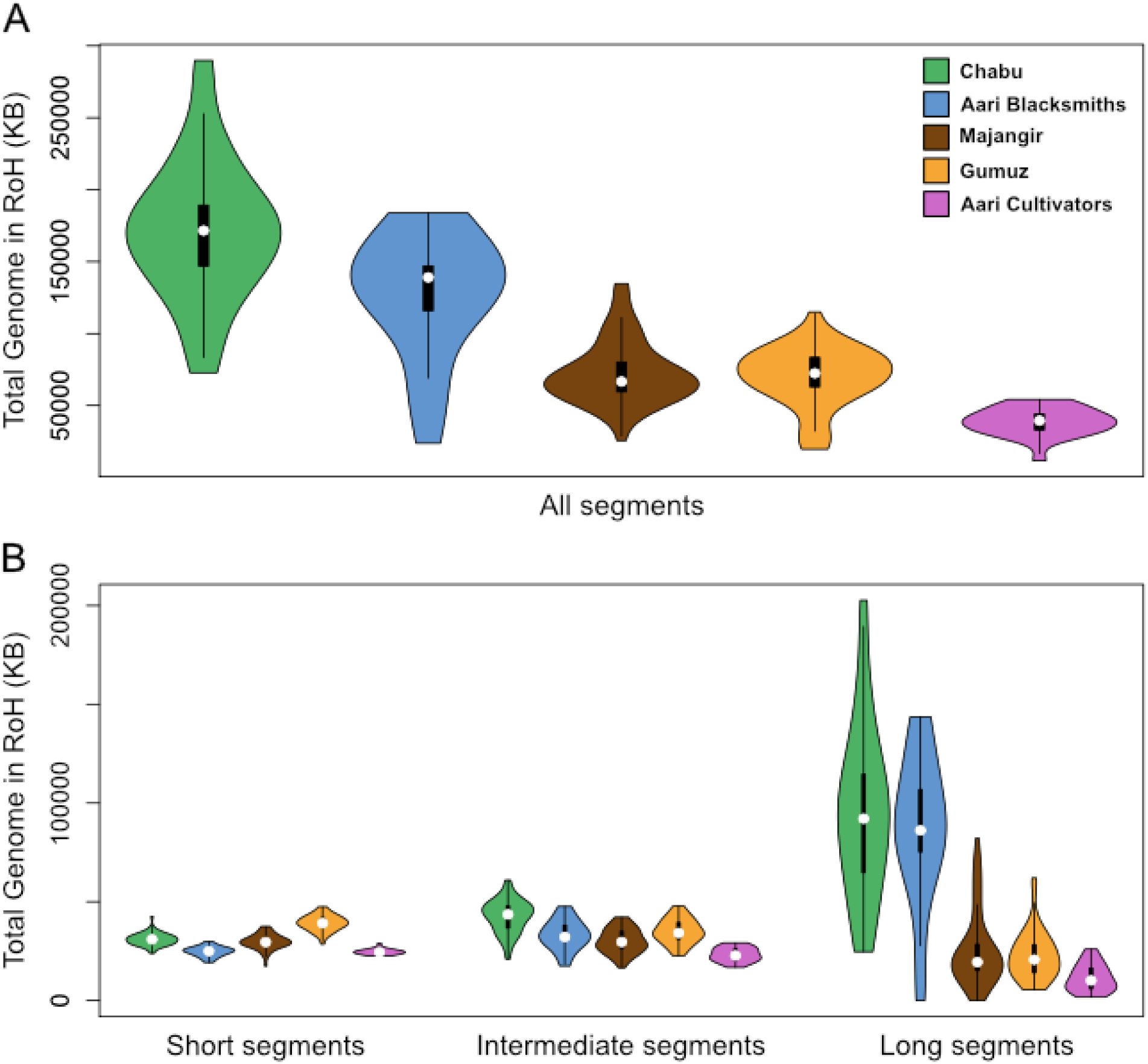
The distributions of the total amount of the genome in runs of homozygosity (RoH) in the Chabu, Aari Blacksmiths, Majangir, Gumuz, and Aari Cultivators for A) all RoH segments and B) in each size class.

Overall, African hunter-gatherer and hunter-gatherer descendant groups vary greatly with respect to total RoH, which suggests that they have experienced different degrees of isolation and/or population size decline in the past. In fact, several hunter-gatherer populations are indistinguishable from farming populations with respect to total RoH (Fig. S5). Interestingly, when using a model-based method of inferring RoH created by ancient, intermediate, and recent events, the Chabu showed significantly elevated total RoH in only the recent (longest) class (Fig. 2). Similar patterns were observed in the Aari Blacksmiths and the Hadza (Fig. 2, S4).

Given this signature of recent demographic pressure on the Chabu, Aari Blacksmiths, and Hadza, we sought to estimate precisely when these population declines occurred. We used a non-parametric method of estimating the effective size (N_e_) of a population through time using the distribution of segments that are shared identical by descent (IBD) across pairs of individuals (*28*). In each population, we optimized the parameters used to infer IBD by comparing RoH, and total the total amount of IBD shared between kin, with ‘truth sets’ generated under independent inference methods (Methods, Figure S6). Through this approach, we estimated historical N_e_ from 4-60 generations ago (ga) and found that the Chabu, Majangir, and Aari Blacksmiths have all faced recent declines in N_e_, while the N_e_ of Aari Cultivators, Gumuz, and Shekkacho have all increased (Fig. 3). The decline in the Chabu and Aari Blacksmiths starting approximately 40 and 50 ga, respectively, is consistent with the RoH results, but the recent Majangir decline starting approximately 50 ga was not suggested by their patterns of RoH. The Biaka, Mbuti, Hadza and Sandawe all have modest estimated N_e_ 60 ga but end up lower by 4 ga, with this decline varying in severity between all four groups. Furthermore, the N_e_ of the Hadza and Mbuti both appear to increase initially (Fig. 3).

**Figure 3.**
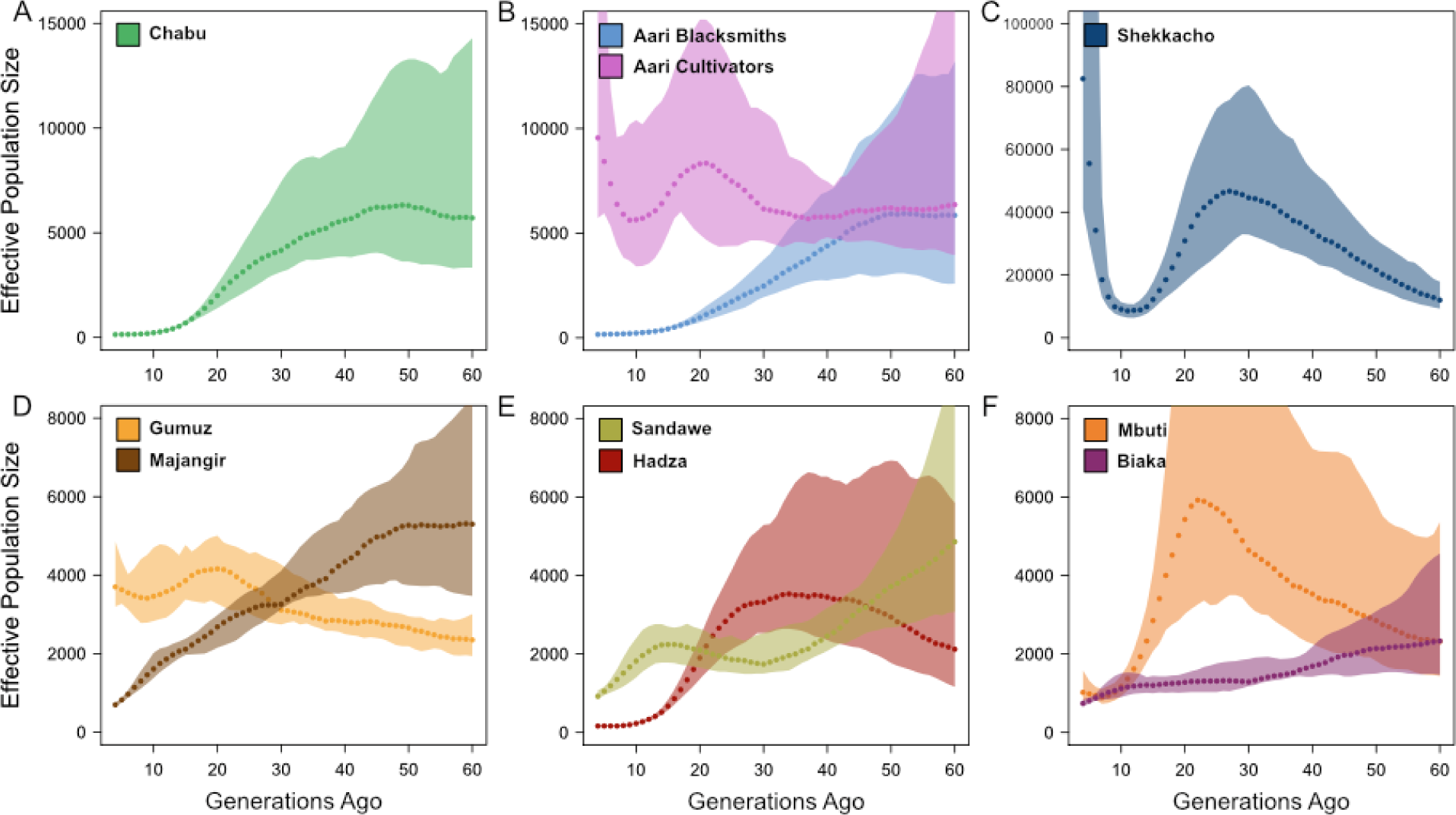
Historical effective population sizes (Ne), from 4 to 60 generations ago, as inferred from distributions of segments that are identical-by-descent across pairs of individuals within A) the Chabu, B), the Aari populations, C) the Shekkacho, D) the Majangir and Gumuz, E) the Sandawe and Hadza, and F) the Mbuti and Biaka. Colored ribbons indicate bootstrapped confidence intervals.

Previous research has shown that European hunter-gatherers that adopted agriculture or were absorbed into agricultural groups experienced population growth (*8*, *29*). Within Africa, the expansion of Bantu-speaking agriculturalists resulted in both the assimilation and extinction of local hunter-gatherers (*9*). By examining a large number of Eastern African groups, our results demonstrate a diversity of hunter-gatherer responses to the intensification and spread of agropastoralism, of which we discuss the five outlined at the beginning of this paper. The Ethiopian Majangir and Gumuz are primarily farmers today, but regularly hunted and gathered in the recent past; both also exhibit many characteristic features of hunter-gatherer societies such as high degrees of egalitarianism and reciprocity (*14*, *25*).

Notably, we show that they have nearly identical ancestry profiles (Fig. 1A, Fig S1). Despite this, their population size trajectories suggest striking historical differences. The Majangir have declined by over 85% from 60 to 4 generations ago, while the Gumuz have increased by more than 50% over the same period. Neither group has seen recent demonstrable gene flow from agropastoralist groups. Similarly, the Sandawe from Tanzania, a ‘click-speaking’ isolate that transitioned to agropastoralism within the past 500 years (*27*), have experienced a pattern of decline that is similar to the Majangir in both trajectory and magnitude. These results may suggest that the ‘late-adopters’, the Majangir and Sandawe, responded to ecological pressure and declining population size by adopting farming with limited gene flow (Response 2).

We also observe opposite demographic trends in the Aari Blacksmiths and Aari Cultivators. Previous studies have shown that these two groups diverged within the last 4,500 years, and are both probable descendants of a ‘Mota-like’ hunter-gatherer population (*21*, *24*). Evidence for a recent bottleneck in the Aari Blacksmiths was also reported (*24*). Our results support these previous findings, and estimate that the decline in the Aari Blacksmiths began approximately 50 ga. We also find that the N_e_ of the Aari Cultivators was relatively steady while that of the Aari Blacksmiths began to decline, and then rapidly increased by half by 4 generations ago. Today, the Aari Blacksmiths are a marginalized group of craftspeople, who are geographically proximate to the Aari Cultivators and Wolayta cultivators, with whom they engage in mutual economic exchange (Response 4) (*24*). Archeological evidence for blacksmithing, today considered a marginal activity across southern Ethiopia (as is foraging and eating wild foods), appears in nearby regions between 1,000 and 3,000 years ago (*30*). It is possible that the genetic divergence of the two Aari populations was associated with the adoption of different cultural practices (i.e. blacksmithing versus farming). These transitions may have influenced the later divergent patterns of historical N_e_ that we observe (Responses 1, 3).

Finally, we show that the Chabu, descendants of earlier Southwestern Ethiopian hunter-gatherers, have experienced a precipitous decline from an ancestral N_e_ of ~5,700 to ~140 over 56 generations. Recent ethnographic work describes the loss of Chabu land to agriculturalists in just the past two decades (*12*); we hypothesize that this is a continuation of a trend that began 40 ga (Response 3). We found that the Chabu and the Aari are descended from Southwestern Ethiopian ancestors that must have once comprised a wide-ranging population (Fig. 1B). Within just the last decade, the Chabu have shown a marked shift in marriage preferences, with an increasing proportion of Chabu men preferring to take a Majangir, Shekkacho, or Amhara spouse (*12*). This may indicate that the Chabu are moving towards assimilating into neighboring agricultural groups, which may lead to further changes in subsistence and culture (Response 1). We found that, similarly to the Chabu, the Hadza of Tanzania underwent a population bottleneck that accelerated over the past 25 generations, reaching a minimum Ne of ~160. The Hadza today live around Lake Eyasi, an area unsuitable for cultivation or pastoralism, which may explain their continued persistence as hunter-gatherers (Responses 3, 5) (*31*).

We characterize nuanced hunter-gatherer responses to recent cultural and demographic changes associated with the spread of agriculture and pastoralism in Eastern Africa. While a shift to agricultural subsistence has been linked to increases in effective population size (*29*), we show a corresponding decline in populations that appear to resist this cultural change. Continued ethnographic and genetic work in collaboration with the Chabu and other marginalized groups will provide valuable insights into the interactions between farmers and hunter-gatherers and the drivers of major cultural transitions.

## Acknowledgments

The authors would like to thank the Stanford Center for Computational, Evolutionary, and Human Genomics and Dr. Carlos Bustamante for providing funding for data generation, and Alexandra Sockell for her assistance with sample preparation. This work was supported by a L.S.B. Leakey Foundation grant to BSH. REWB was supported by an Exploration Fund grant from The Explorers Club, the NSF IGERT Program in Evolutionary Modeling at Washington State University, and the Max Planck Institute for the Science of Human History.

## Ethics statement

REWB collected the Chabu samples after months of ethnographic research with the communities by Samuel Dira (SD) and BSH. The research with the Chabu and their neighbors is part of a larger formal collaborative research and capacity building relationship between the Departments of Anthropology at Hawassa University, Ethiopia (HU) and Washington State University (WSU). The collaboration involves training several HU faculty, such as SD, in the WSU PhD program and cooperative participation in research projects in Southwestern Ethiopia. Authorization for the research was obtained from HU and WSU.

## Materials and Methods

### Collection, Consent, Ethics approval

Samples from the Chabu, Majangir, and Shekkacho were collected by REWB in May 2013, using Oragene•DISCOVER (OGR-500) kits for the Chabu and generic 5 ml tubes with Norgen preservation solution for the other two groups. Prior approvals for the project were obtained from the leadership of each group, from the School of Behavioral Sciences at Hawassa University (#BS/502/05), and from the Majangir Zone Council of the Gambella Regional State (#901/ማዞ1/ፌ4). Ethical approval for human subjects research was obtained from the Institutional Review Board of Washington State University under proposals #12972 and #13134. Informed consent was obtained from each participant after reading or hearing the approved text translated into their local language and providing their signature, or a fingerprint in lieu of a signature for non-literate participants.

### Data generation and processing

50 individuals each of the Chabu, Majangir, and Shekkacho were genotyped using the Illumina Infinium MultiEthnic Global Array, which assays over 1.7 million genetic markers. Genotypes were initially called using Illumina GenomeStudio software. We removed 5 samples (1 Majangir, 4 Shekkacho) that had a call rate below 90%. Calls for rare variants, defined as those having a minor allele frequency (MAF) < 5%, were then replaced by using zCall following their published procedure (*32*). Variants with more than 15% missing data, an observed heterozygosity greater than or equal to 80%, or with cluster separation less than or equal to 2% were removed from the dataset. In preparation for merging with additional datasets, we converted all variants to the Illumina top strand and oriented them to match the 1000 Genomes reference. We renamed SNPs to match dbSNP version 144, and removed all indels and A/T or C/G transversion variants, leaving over 1.3 million SNPs in the final dataset.

### Unsupervised clustering and principal components analyses of genotype data

We merged our Ethiopian SNP data with previously published or publically available genotype data from other Ethiopian, Somali, Sudanese, South Sudanese, as well as the 1000 Genomes Yoruba, Luhya, Maasai, and Iberians (*33*–*36*). We removed SNPs that did not overlap across datasets, were out of Hardy-Weinberg equilibrium (p < 0.001) in any population, or had a missingness rate of over 5% in the merged dataset. We also removed SNPs with a MAF of 5% or less and SNPs in linkage disequilibrium (r^2^ > 0.1 within a 50 kbp window, stepping 10 kbp at a time), leaving 59,700 SNPs and 1233 individuals for analysis across 37 populations, plus Mota. Using this dataset, we identified and removed related individuals within populations using kinship statistics calculated by PLINK (PI_HAT > 0.1). Of the remaining individuals, we randomly discarded a set of individuals such that no population, as defined by our population labels, had more than 50 individuals. We then applied a less strict linkage disequilibrium filter (r^2^ > 0.3 within a 50 kbp window, stepping 10 kbp at a time) to the MAF filtered data, leaving a dataset of 106,997 SNPs from 828 individuals across 37 populations plus Mota. We ran the ADMIXTURE algorithm for Ks of 2 to 12 with 10 replicates each (*37*). We used *pong* to visualize the concordance between different runs and, for each K, identify the most frequent mode among the 10 replicates (*38*). For extant populations with known sampling or ethnographic coordinates (see ‘Estimated effective migration surfaces’), we also plotted the population averages of each ancestry component geographically, interpolating between datapoints across the landscape as in Uren et al. 2016 (*39*). We also used the merged and LD filtered genotype dataset to perform a principal components analysis (PCA) using *smartpca* (*40*).

### Estimated effective migration surfaces

We used a method of estimating effective migration surfaces (EEMS) to visualize variation in migration rates across East Africa (*22*). The algorithm takes geo-referenced genetic SNP data as input and simulates migration across a grid under a stepping-stone model, returning a spatial depiction of estimated historical rates of gene flow. We prepared the SNP dataset by following the same procedure as for ADMIXTURE and PCA, but excluded some samples before relatedness and LD filtering. The samples we removed were Mota, the ancient sample, the Somali and Sudanese populations from Pagani et al. 2012, due to lack of specific geographical information, and the Iberians and Yoruba because they live outside of the region of interest. The Baria were excluded in this filtering process for having excessive missing data. This left a dataset of 131,892 SNPs and 571 individuals across 32 populations.

We primarily used the coordinates in the original publication with some adjustments. As this is a spatial analysis based on historical population locations, the coordinates for Tigray individuals were changed from their sampling location near Addis Ababa to their traditional homeland in Eritrea, using the Glottolog coordinates for Tigrinya (*41*). For similar reasons, we excluded the 3 Hausa individuals given their recent migration from outside the region of interest within the last 100 years (*42*). We found that neither of these changes led to major qualitative differences in results. We performed a number of runs under a range of starting parameters (number of demes specified as 200, 300, 400, and 500, each under three different starting seed values) and averaged the results to mitigate the possible bias of any single run. Each run was allowed to proceed for 30 million MCMC iterations to ensure convergence, with the first 15 million discarded as burn-in and the remaining 15 million thinned to retain 1 out of every 15,000 data points. Proposal variances were tuned so that proposals were accepted between 20% and 30% of the time for all runs.

### Estimating the distributions of major language families

In order to determine the correspondence between EEMS-inferred migration barriers and corridors and linguistic boundaries, we calculated kernel estimates of language family distributions using the adaptive radius local convex hull (*a*-LoCoH) method (*43*). Language centroid point data and (Greenberg-based) family classifications for every known living African language were obtained from Ethnologue (https://www.ethnologue.com/). We then applied the *a*-LoCoH algorithm to construct ‘utilization distributions’ for each of the five major African language families (Niger-Congo, Afro-Asiatic, Nilo-Saharan, Khoisan, and Austronesian), using values of *a* equal to the longest geodesic distance between any two languages in a family, to accommodate variable point densities. This produced a set of layered isopleths for each language family representing decile occurrence probabilities. These isopleths were then plotted with overlaid language point data to visualize the extent and density of language distributions by family in relation to historical migration rates estimates as determined by EEMS (Fig. S2C). Putative linguistic isolates were determined according to Blench, 2017 (*44*).

### Runs of homozygosity

We determined runs of homozygosity (RoH) in the autosomes of the Ethiopian populations and other African hunter-gatherers, the Hadza, Sandawe, Biaka, and Mbuti. We used the GenomeStudio and zCall processed dataset for the Chabu, Majangir, and Shekkacho, a merged dataset of Ethiopians assayed on the Illumina Infinium Omni 1M and Omni 2.5M arrays (*33*, *34*), a merged dataset of Southern African hunter-gatherers and Western African hunter-gatherers assayed on the Illumina 550k and 660k arrays, respectively (*26*, *36*). We removed SNPs with more than 5% missingness or a less than 1% MAF from all datasets. We also randomly thinned the Ethiopian datasets to approximately match the SNP density of the merged Southern and Western African hunter-gatherer dataset. Lastly, we removed SNPs that were not in Hardy-Weinberg equilibrium within each population (p < 0.001), which left approximately 470,000 SNPs per population for analysis. We then identified RoH in each individual using PLINK, defining a run as having at least 50 SNPs and being at least 1 Mb in length, allowing for no more than two missing and one heterozygous SNP per run. Despite varying these parameters, we found many instances of two RoH within a single individual closely flanking a low SNP density region. We chose to join such segments post hoc with a custom script by defining low density regions as 1Mb windows that fell in the lower 5% of SNP count when compared to the entire genome. We also observed genome regions where unusually high numbers of individuals in a population carried a RoH segment. We defined such outlier SNPs as being more than three standard deviations above the mean depth of RoH in the population. We joined nearby outlier SNPs into larger regions and added them to a list of previously identified low density regions, known low complexity regions (i.e. heterochromatin, telomere, centromere, and short arm regions). We removed all RoH segments that overlapped by 85% or more with one of these regions.

### GARLIC

In order to analyze RoH in separate classes corresponding to the age of the events that produced them, we also used GARLIC software to identify RoH in each population (*45*). This algorithm implements a population model-based method of inferring RoH in ‘short’, ‘intermediate’ and ‘long’ size classes (*46*). We ran GARLIC on the missingness- and MAF-filtered datasets used for PLINK RoH analysis, after removing SNPs that were out of Hardy-Weinberg equilibrium within the population (p < 0.001), using the following parameters: ‘error’ of 0.001, ‘winsize’ of 30, ‘auto-winsize’, and ‘auto-winsize-step’ of 5. As with PLINK RoH, we then joined segments that flanked regions of low SNP density, and updated their size class accordingly.

### IBDNe

Working from the missingness- and MAF-filtered datasets used for PLINK RoH analysis, we phased all individuals in a given dataset together using SHAPEIT2 with a window size of 5 Mb and the duoHMM option and haplotypes from the 1000 Genomes phase 3 dataset as a reference. We converted the output of SHAPEIT2 to a ‘phased’ PLINK file format using a custom script. We then used GERMLINE2 to identify tracts shared identical-by-descent (IBD) between chromosomes across all individuals in a given dataset using the --w_extend and --haploid flags (*47*). We then joined IBD segments that, perhaps due to errors in phasing or genotyping, were separated by a gap of less than 0.6 centiMorgans that contained no more than one discordant SNP (*28*). As described in the ‘Runs of homozygosity’ section, we also filtered out segments that had high overlap with regions of excess depth, low complexity, or low SNP density. In inferring IBD tracts, we varied the ‘bits’ (from 5 to 200, increasing by 5) and ‘errhom’ (between 0, 1 and 2) parameters and decided the optimal combination for each population using three metrics. These parameters control the minimum number of exactly matching SNPs required to call an IBD segment and the number of mismatches allowed, respectively (*47*). First, we compared the distribution of RoH inferred by GERMLINE2 to that inferred by PLINK, and calculated how much of the RoH inferred by GERMLINE2 was not inferred by PLINK, normalizing by the total amount of RoH inferred by PLINK. Second, we calculated how much of the RoH inferred by PLINK was not inferred by GERMLINE2, normalizing by the total amount of RoH inferred by PLINK. Finally, we compared the total amount of the genome inferred to be IBD between pairs independently by PLINK and GERMLINE2, and calculated the residuals. We selected the GERMLINE2 parameter combination that produced the IBD segment distribution that most closely represented the PLINK results based on the average of these three metrics (Table S1). We then used IBDNe to estimate the historical effective population size from all IBD segments shared within a population that were 4 centiMorgans or longer (*28*).

### Political boundaries in maps

The boundaries depicted in the maps do not imply the expression of an opinion by any of the authors of this paper regarding the legal status or political boundaries of any country or territory.

---

**Figure S1.**
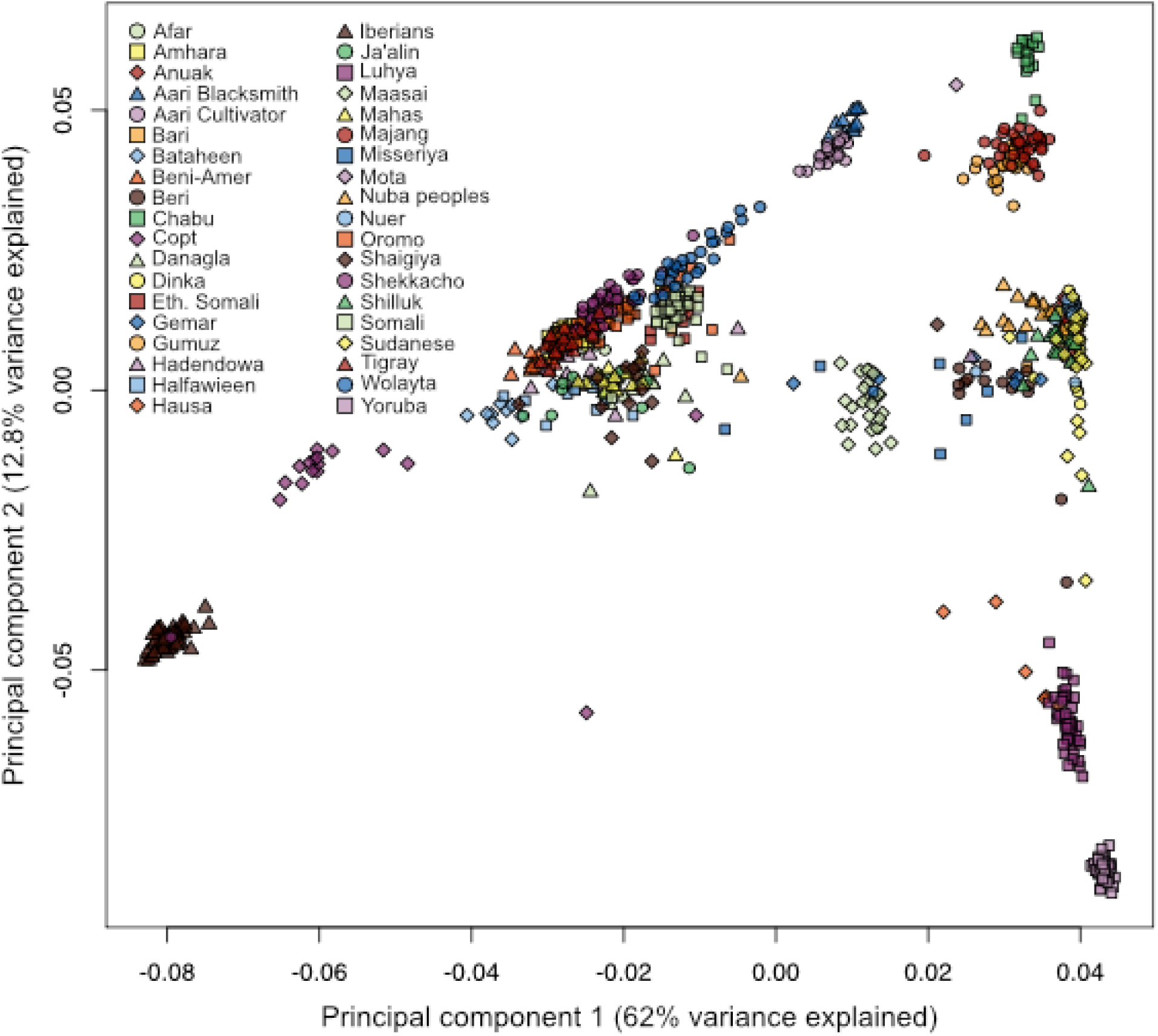
Principal components analysis on genotype data from populations of northeast Africa, including Mota, an ancient individual, as well as the Yoruba from Nigeria and Iberians from Spain.

**Figure S2.**
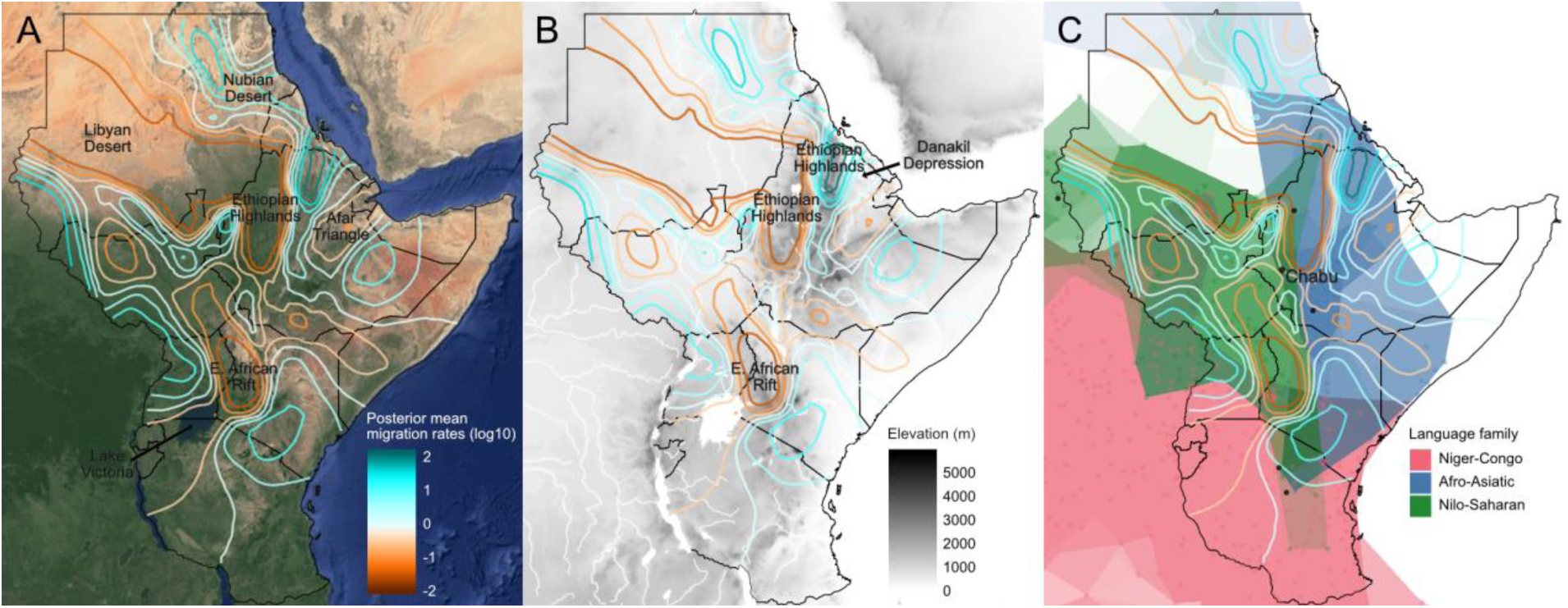
Effective migration surfaces depicted as contour lines over A) satellite imagery, B) elevation and water features, and C) the geographic distribution of major language families in Eastern Africa (see Methods). For contour lines, cool colors correspond to effective migration corridors, while warm colors correspond to effective migration barriers. A) Some geographic features, such as the Libyan Desert (Sahara) in northwestern Sudan, the northwestern Ethiopian Highlands, the Afar Triangle in the Ethiopian Rift down through the East African Rift, and Nalubaale/Lolwe/Nyanza (Lake Victoria), correspond with historical migration barriers. Map data from Google; imagery © 2018 TerraMetrics. B) Some regions of high elevation or roughness, such as the northwestern Ethiopian Highlands and the volcanic range along the East African Rift, correspond with historical migration barriers. However, the northeastern Ethiopian Highlands feature high estimated rates of historical migration, potentially as a preferred route around the inhospitable Danakil Depression to the east. Elevation data at 30 arc-second (~1 km) resolution are from the U.S. Geological Survey GOTOPO30 digital elevation model, accessed through EarthExplorer (https://earthexplorer.usgs.gov/). Physical vectors for rivers and lakes at 1:10 million scale are from Natural Earth (https://www.naturalearthdata.com/). C) Darker isopleths represent higher occurrence probabilities for languages within each family. Putative linguistic isolates, including the Chabu, are depicted as black points. Language data are from Ethnologue (https://www.ethnologue.com/).

**Figure S3.**
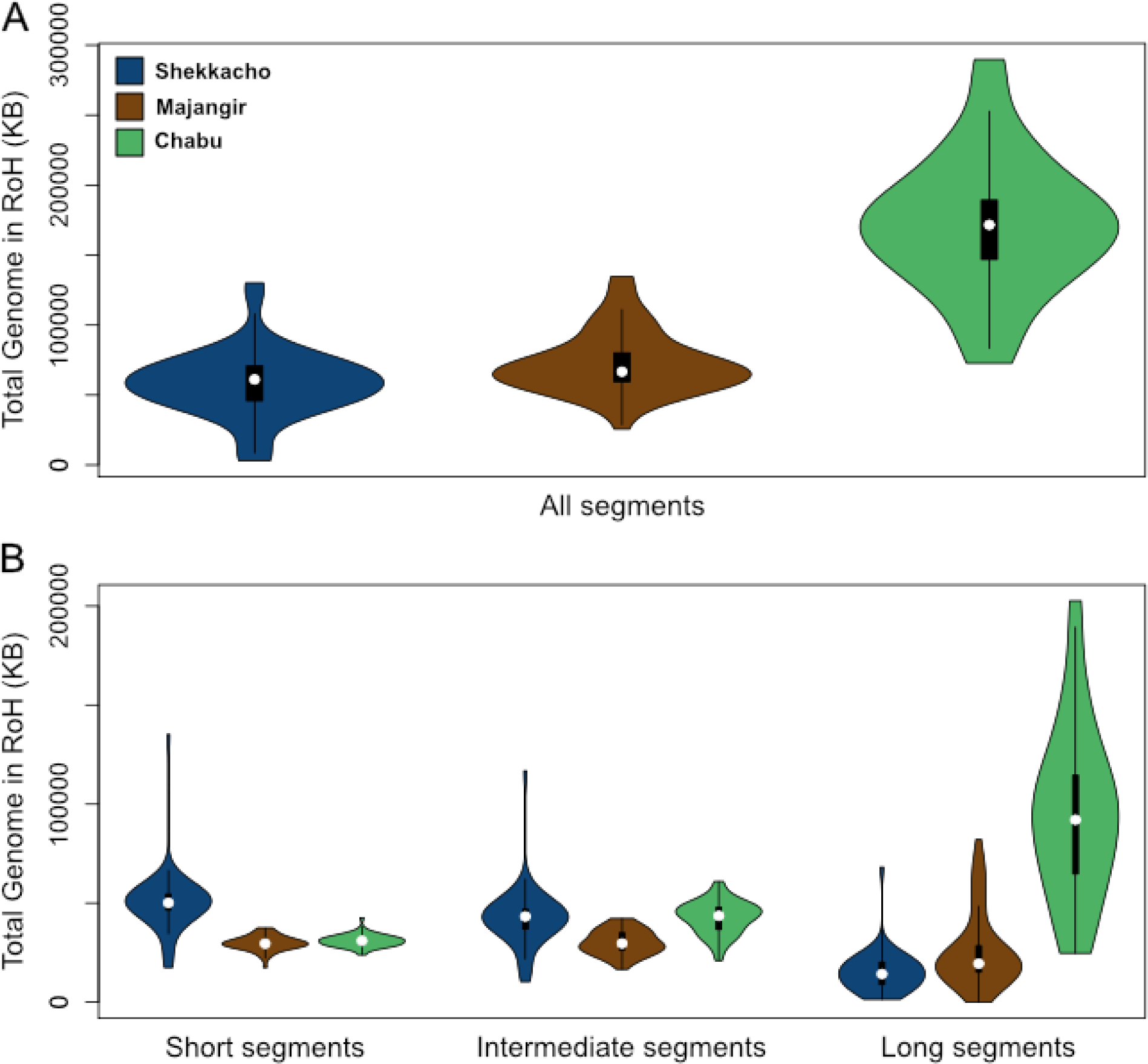
The distributions of the total amount of the genome in runs of homozygosity (RoH) in the Shekkacho, Majangir, and Chabu for A) all RoH segments and B) in each size class.

**Figure S4.**
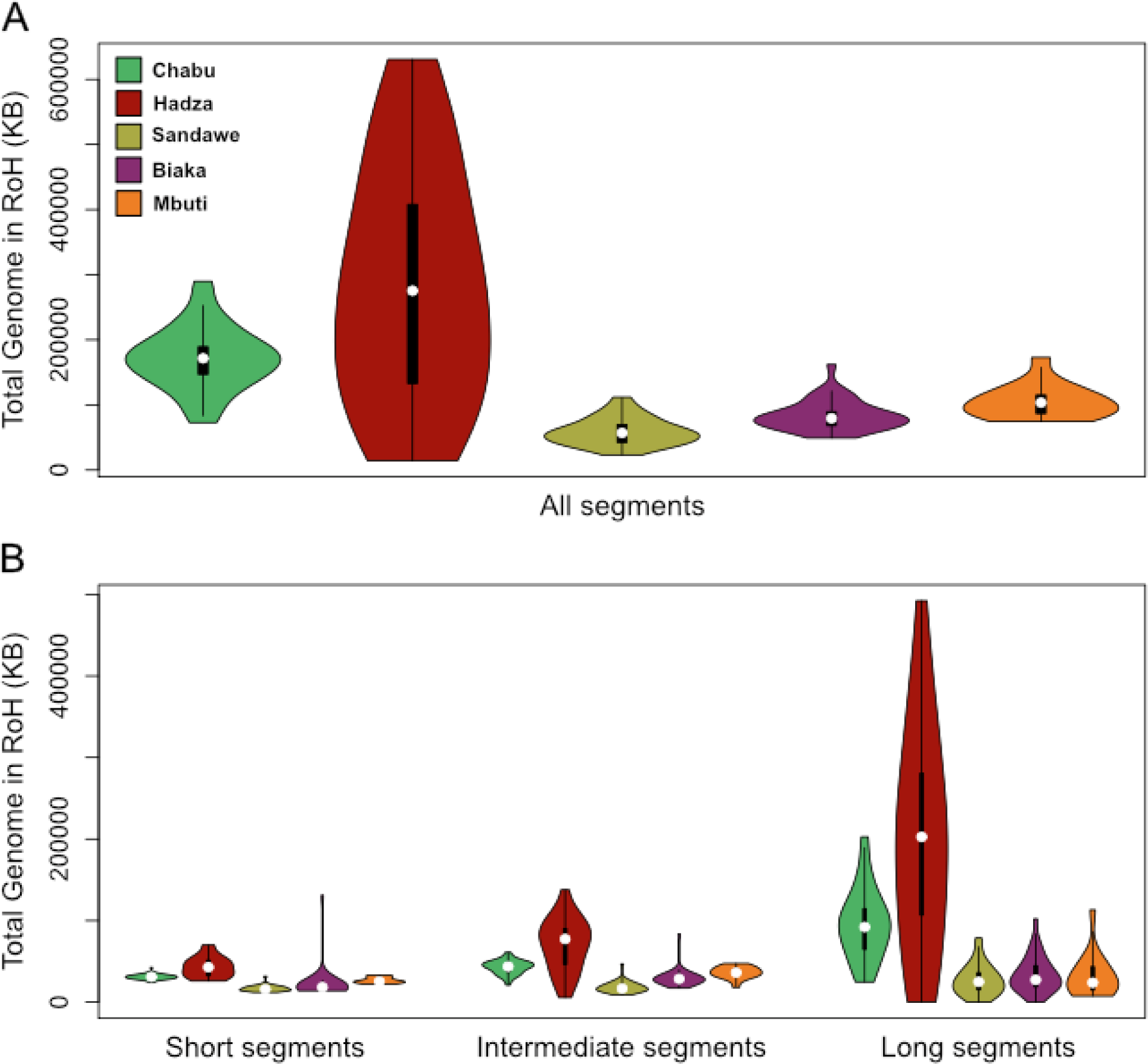
The distributions of the total amount of the genome in runs of homozygosity (RoH) in the Chabu, Hadza, Sandawe, Biaka, and Mbuti for A) all RoH segments and B) in each size class.

**Figure S5.**
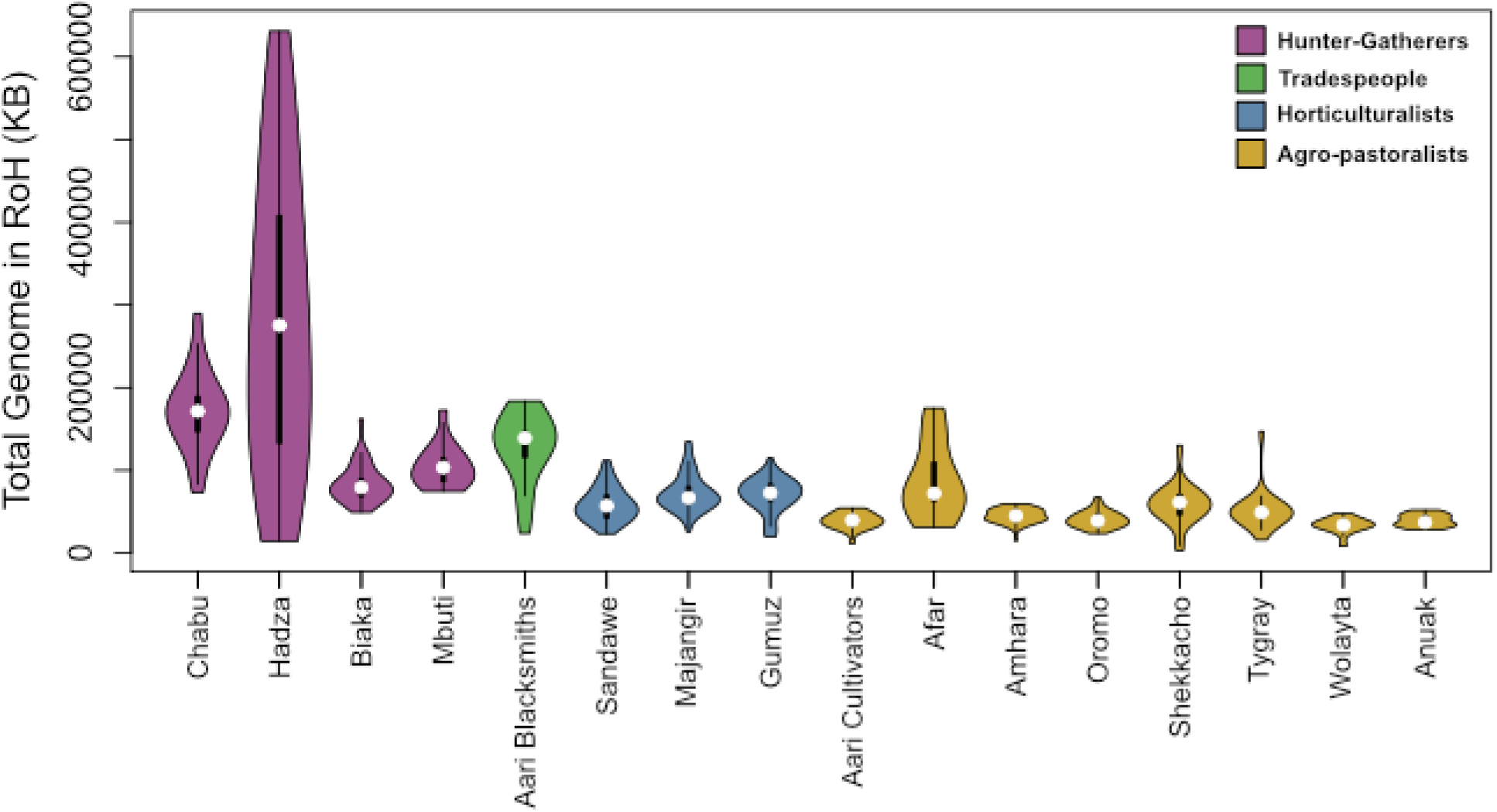
The distributions of the total amount of the genome in runs of homozygosity (RoH) in all Ethiopian populations and African hunter-gatherers. Populations are grouped and colored according to their primary subsistence or economic strategy.

**Figure S6.**
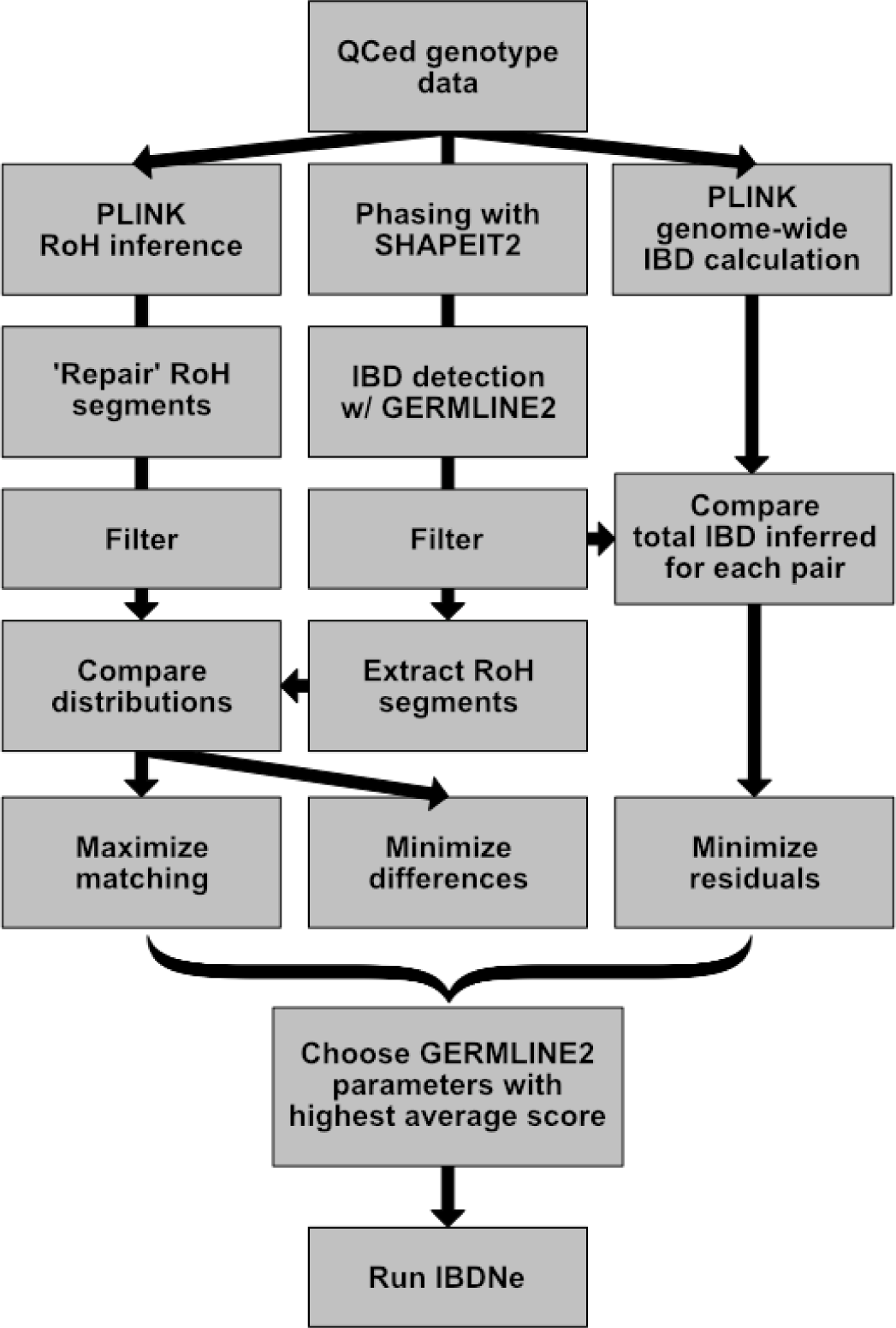
A schematic of the strategy used to select the optimal parameters for GERMLINE2 to infer the IBD segment distribution for each population.

**Table S1.**
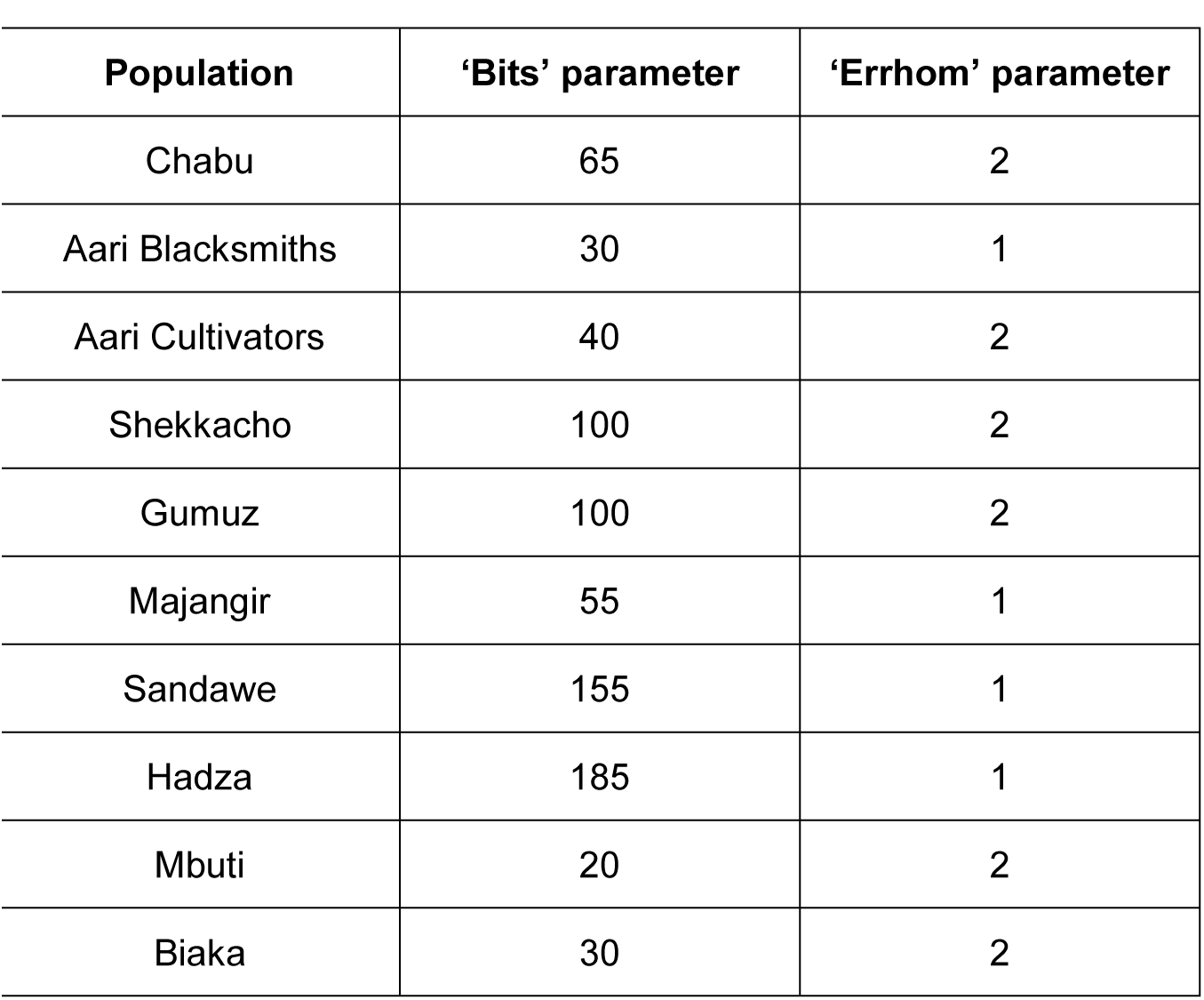
The GERMLINE2 parameters chosen for each population by our optimization strategy for the inference of historical effective population size (N_e_) (see Materials and Methods).

